# Prediction of local convergent shifts in evolutionary rates with *phyloConverge* characterizes the phenotypic associations and modularity of regulatory elements

**DOI:** 10.1101/2022.05.02.490345

**Authors:** Elysia Saputra, Weiguang Mao, Nathan Clark, Maria Chikina

**Affiliations:** Joint Carnegie Mellon University – University of Pittsburgh PhD Program in Computational Biology, Pittsburgh, PA, USA; Department of Computational and Systems Biology, University of Pittsburgh, Pittsburgh, PA, USA; Department of Human Genetics, University of Utah, Salt Lake City, UT, USA; Pittsburgh Center for Evolutionary Biology and Medicine, University of Pittsburgh, Pittsburgh, PA, USA

## Abstract

Physiological and morphological adaptations to extreme environments arise from the molecular evolution of protein-coding regions and regulatory elements (REs) that regulate gene expression. Comparative genomics methods can characterize genetic elements that underlie the organism-level adaptations, but convergence analyses of REs are often limited by their evolutionary properties. A RE can be modularly composed of multiple transcription factor binding sites (TFBS) that may each experience different evolutionary pressures. The modular composition and rapid turnover of TFBS also enables a compensatory mechanism among nearby TFBS that allows for weaker sequence conservation/divergence than intuitively expected. Here, we introduce *phyloConverge*, a comparative genomics method that can perform fast, fine-grained local convergence analysis of genetic elements. *phyloConverge* calibrates for local shifts in evolutionary rates using a combination of maximum likelihood-based estimation of nucleotide substitution rates and phylogenetic permutation tests. Using the classical convergence case of mammalian adaptation to subterranean environments, we validate that *phyloConverge* identifies rate-accelerated conserved non-coding elements (CNEs) that are strongly correlated with ocular tissues, with improved specificity compared to competing methods. We use *phyloConverge* to perform TFBS-scale and nucleotide-scale scoring to dissect each CNE into subregions with uneven convergence signals and demonstrate its utility for understanding the modularity and pleiotropy of REs. Subterranean-accelerated regions are also enriched for molecular pathways and TFBS motifs associated with neuronal phenotypes, suggesting that subterranean eye degeneration may coincide with a remodeling of the nervous system. *phyloConverge* offers a rapid and accurate approach for understanding the evolution and modularity of regulatory elements underlying phenotypic adaptation.

## Introduction

Decoding the genetic basis of complex phenotypes is a central goal of biology, and one strategy for learning genotype-to-phenotype associations is by studying the genetic basis of morphological adaptation. When species transition to a new environment, accompanying shifts in selection pressures can cause numerous molecular changes that give rise to phenotypic alterations at the organismal level. Morphological and physiological adaptations are enabled by changes in both protein-coding elements and regulatory elements that play key roles in determining gene expression patterns in different contexts (Wray 2007; Carroll 2008).

With the wealth of sequenced species genomes that has been produced by high-throughput sequencing, it is possible to identify the functional associations of genetic elements by comparing the sequences of species with an extreme phenotype with orthologous sequences in other species. Convergent evolution is a useful phenomenon that allows us to distinguish phenotype-associated evolutionary processes from lineage- or species-specific changes that cannot be attributed to specific selection pressure. When independent lineages convergently adapt to a common selection pressure, genetic elements that control the selected phenotypes are likely to undergo similar selective shifts. Some genetic elements that experience stronger selective constraints would shift to a slower evolutionary rate, while other genetic elements, such as those supporting functionality no longer needed in the new environment, may experience relaxed constraints and accumulate more divergence. This relationship between selection and sequence conservation has given rise to parameter models for detecting lineage-specific rate shifts (Pollard et al. 2006; Siepel et al. 2006), as a successive step toward convergent rate shifts across disjoint clades. The relationship between convergent phenotypes and convergent rate shifts can be exploited to associate genetic elements with high-level phenotypic adaptations. The utility of this comparative framework has been successfully demonstrated in numerous studies (Prudent et al. 2016; Partha et al. 2017; Meyer et al. 2018; Roscito et al. 2018; Hu et al. 2019; Kowalczyk et al. 2020) and engendered several computational algorithms (Hiller et al. 2012; Marcovitz et al. 2016; Prudent et al. 2016; Hu et al. 2019; Kowalczyk et al. 2019). Of the existing methods, Forward Genomics (Hiller et al. 2012; Prudent et al. 2016) and RERconverge (Kowalczyk et al. 2019; Partha et al. 2019) stand out as having been applied at genome-wide scale to a variety of different phenotypes.

The methods have demonstrated success in identifying genome-wide phenotypic associations for both protein-coding and non-coding elements, but their application to non-coding regions is limited because such methods require a defined unit of non-coding sequence to operate on. The typical strategy for defining non-coding units is to use PhastCons (Siepel et al. 2005), which segments the alignment into conserved regions. This approach produces a set of conserved non-coding elements (CNEs) that represent putative regulatory elements (REs) and have a size range of 50-500bp, much larger than a single transcription factor binding site (TFBS). This disconnect between the CNE unit and the TFBS, which is the atomic unit of sequence activity, poses specific challenges for evolutionary analysis.

REs typically contain multiple TFBS for different TFs (though often with some repetition) (Long et al. 2016). Detailed experiments on dissection of well-characterized REs have revealed that the relationship between individual TFBS and the functional output of the RE is complex. Ablating TFBSs may eliminate activity, change it, or have no effect (Patwardhan et al. 2012; Preger-Ben Noon et al. 2018; Serrano-Saiz et al. 2020; Wong et al. 2020; Jindal and Farley 2021; Snetkova et al. 2021). Moreover, RE activity is itself multifactorial as many REs are pleiotropic and can drive expression in seemingly unrelated contexts. These pleiotropic effects can occur via identical TFs binding to identical sites, different TFs binding to identical sites, and different site usage. All three scenarios have been observed (Spivakov 2014). From the perspective of genome-wide evolutionary analysis of CNE (which are computationally identified putative REs), individual TFBSs may have different and possibly context-specific contributions to regulatory activity and thus have different evolutionary pressure and histories. It is thus quite likely that phenotype-driven changes in evolutionary rate may be more localized than the typical CNE length. As such, the information content across a given CNE may be non-uniform, making it necessary to interrogate a genetic element at a higher resolution. There is therefore a need for a computational strategy that allows us to scan a multiple sequence alignment (MSA) and identify functional units of REs without prior information on the defined units of non-coding elements.

To address these challenges, we present *phyloConverge*, a fast comparative genomics method that performs fine-grained local convergence analysis to identify genomic regions associated with phenotypic convergence at high resolution. Our method combines explicit parameterization of evolutionary rate shifts and a phylogeny-aware trait permutation strategy to produce unbiased conservation/acceleration scores calibrated to the local context of the chromosomal region. By requiring a single phylogenetic tree estimation step and employing an adaptive strategy for accelerating permutation tests, *phyloConverge* can perform rapid high-resolution scanning of individual nucleotides across a MSA to detect convergent shifts in evolutionary rates correlated with convergent phenotypic evolution.

## Results

### The *phyloConverge* method augments *phyloP* functionality with methods to correct for biases in convergent trait analysis

Given a MSA of a region (or a nucleotide position) of interest, a phylogenetic model of neutral nucleotide substitution, and a defined set of convergent species (i.e., “foregrounds”), *phyloConverge* measures evolutionary rate shifts that occur across the foreground branches while correcting for statistical bias. *phyloConverge* builds on the widely used *phyloP* framework (Pollard et al. 2010) to detect rate shifts that convergently occur on the foreground branches relative to the remaining branches in the phylogeny (i.e., “backgrounds”). A convergent rate shift is inferred by performing maximum likelihood estimation of two branch scaling factors (Fig. 1A). The first scaling factor *ρ*, measuring the phylogeny-wide rate shift relative to the provided neutral tree, is analogous to the parameter used to compute the widely used *phyloP* conservation track. An additional *λ* parameter measures evolutionary rate shifts that occur exclusively among the foregrounds relative to the entire phylogeny.

**Figure 1.**
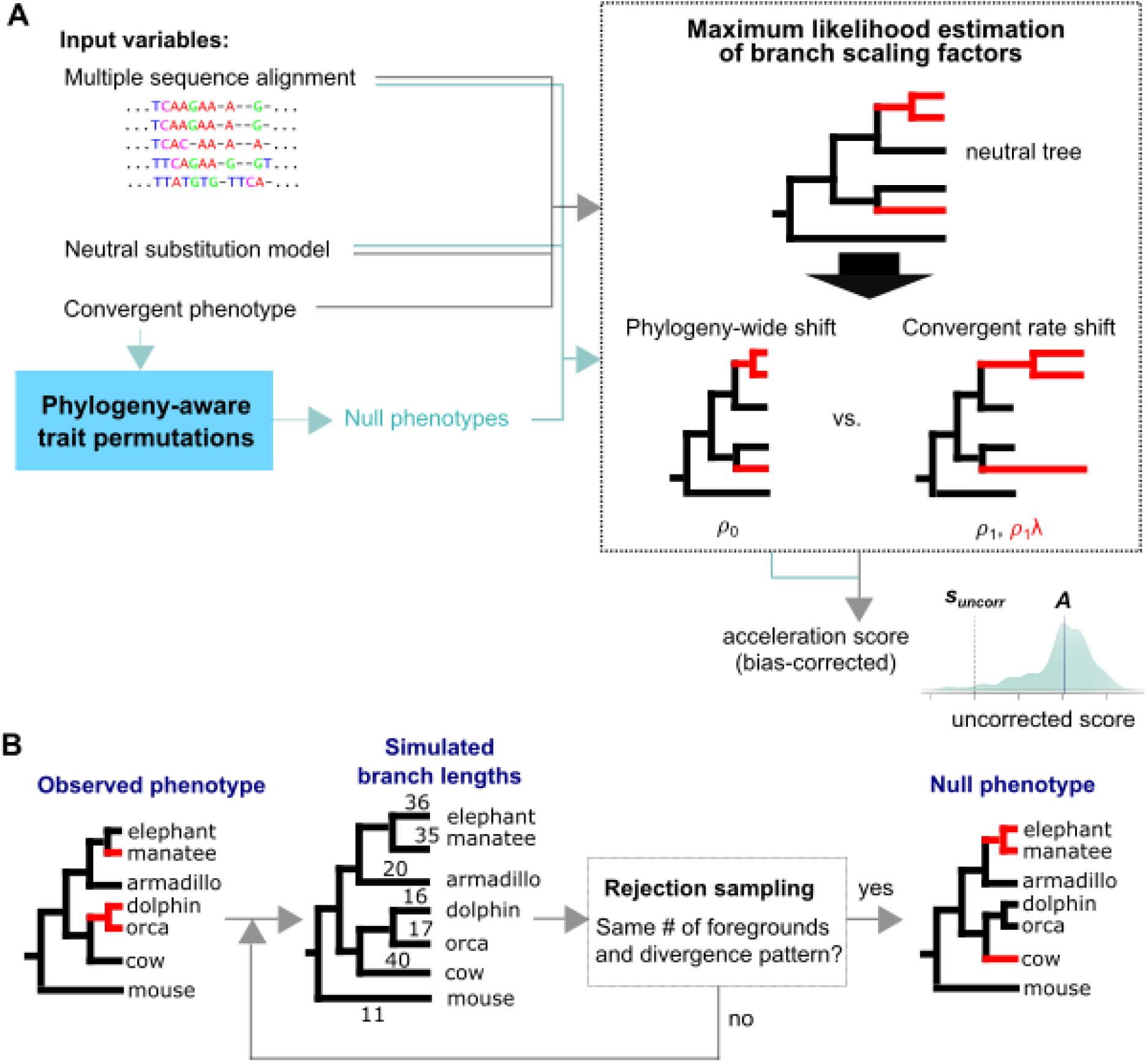
Workflow of *phyloConverge*. (**A**) Given input variables that include the set of species with convergent phenotype (“foreground” species), the neutral model of evolution, and the multiple sequence alignment, *phyloConverge* combines generative nucleotide substitution modeling and phylogeny-aware trait permutation to compute rate acceleration/deceleration scores that are empirically corrected for statistical biases. Using maximum likelihood estimation of branch scaling factors (*ρ_o_* and *ρ_1_* denoting phylogeny-wide scaling factors in *H_o_* and *H_1_*, respectively, and *λ* denoting foreground-specific scaling factor), *phyloConverge* performs likelihood ratio test to test the null hypothesis that the foreground branches are not differentially scaled from the background branches, and then conducts this test on numerous “fake” phenotypes to correct for biases. *phyloConverge* computes conditional two-tailed p-values that accounts for non-trivial distributions of uncorrected scores, given a defined centerpoint *A* of the distribution (set as the median). (**B**) *phyloConverge* uses phylogenetic permulation to produce null phenotypes that preserve the covariates in the observed convergent phenotype. *phyloConverge* performs phylogenetic simulations to produce simulated branch lengths and select the top-ranking branches as candidates for a new fake phenotype. These candidate null foregrounds are accepted if they match the observed foregrounds in terms of number and phylogenetic dependence.

Given these definitions, evidence for rate convergence is quantified by a likelihood ratio test (LRT) comparing the null hypothesis of constant scaling across both foreground and background (i.e., all branches are uniformly scaled by *ρ_o_*) against the alternative hypothesis that the foreground branches are scaled by *λ*, in addition to the background scaling *ρ_1_*. After estimating these parameters and performing hypothesis testing, the conservation/acceleration score is finally defined by computing the negative log-likelihood of the LRT p-value, noting the magnitude of *λ_1_* (conservation if *λ_1_* < 1, acceleration if *λ_1_* > 1).

*phyloConverge* then incorporates a phylogeny-aware empirical bias correction to account for statistical biases that have not been effectively captured by this two-parameter model. It has been previously noted that phylogenetic inference methods can produce highly skewed statistics when testing the same hypothesis across a large collection of genetic elements. This phenomenon is not specific to particular inference methods, and indeed even occurs in the context of phylogenetic generalized least squares (pGLS), but is a general problem that arises from the hypothesis having shared bias structure (Hadfield and Nakagawa 2010; Stone et al. 2011; Sakamoto and Venditti 2018; Kowalczyk et al. 2020; Saputra et al. 2021). Such biases can arise from failing to completely account for phylogenetic dependence (Felsenstein 1985; Engstrom et al. 2007; Touceda-Suárez et al. 2020) or systematic variation across genomes either of biological (variation in nucleotide content (Romiguier and Roux 2017)) or technical (genome quality (Hosner et al. 2016)) origin. When testing a single hypothesis genome-wide, these subtle effects induce test dependency that results in highly skewed p-value distributions. Furthermore, in the case of *phyloConverge*, the LRT is not even expected to produce well behaved p-values for the simple reason that the single parameter scaling model is an oversimplification. In reality, there is considerable variation in evolutionary rates across the genome. Thus, when scaling with respect to a single neutral tree, the most accurate model would give each branch its own scaling parameter to account for local variation. Consequently, increasing the number of parameters from one to two will often produce a significantly better fit even in the absence of a specific foreground signal.

To correct for these biases, we previously developed a phylogenetic trait permutation method called permulation, a portmanteau of *permu*tation and simu*lation* (Saputra et al. 2021). Permulation is a rejection sampling approach that uses Brownian motion simulations to produce multiple “fake” (null) traits by selecting new sets of foreground species that are matched to the true observed trait in terms of number of species and phylogenetic dependence (Fig. 1B). With these null traits, we can perform the equivalent of permutation tests to correct the test statistics. We incorporate this trait permulation strategy into *phyloConverge* to produce *n* null traits and use *phyloP* to compute the conservation/acceleration scores for both the observed convergent trait and the set of *n* null traits. Finally, we measure the corrected significance of rate shift by computing an empirical p-value *p_corr_*, defined as the proportion of the null *phyloP* scores that are as extreme or more extreme than the *phyloP* score of the true phenotype. The corrected conservation/acceleration score *s_corr_* is then defined as the negative logarithm of *p_corr_*, signed by the direction of rate shift (deceleration or acceleration). Specifically, *s_corr_* > 0 denotes stronger foreground conservation, whereas *s_corr_* < 0 denotes foreground acceleration (see Methods for details).

While such a permutation test is important for accurately calibrating our confidence in the identified genotype-phenotype associations, the main drawback is that it necessitates a large number of computations to achieve a high p-value resolution, which increases running time significantly. To overcome this drawback, we adopted an adaptive permutation strategy previously applied in expression quantitative trait loci (eQTL) analysis (Wang et al. 2020), which balances p-value resolution against running time by pruning the number of permutations if a certain significance threshold has been crossed and computing an adaptive p-value (see Methods).

### *phyloConverge* improves detection of elements underlying convergent phenotypes

We first benchmark *phyloConverge* using a well-characterized convergent trait, the subterranean mammal habitat. We use a dataset that was previously analyzed by Roscito et al. (Roscito et al. 2018), which contains a MSA of 24 species including 4 subterranean mammal lineages: the naked mole rat, the cape golden mole, the star-nosed mole, and the blind mole rat (Fig. 2A). Using the PhastCons tool (Siepel et al. 2005), Roscito et al. had previously computed 491,576 conserved non-coding elements (CNEs) that align well among at least 15 species in the phylogeny and used Forward Genomics (Prudent et al. 2016) to identify 9,364 “subterranean-accelerated” CNEs. Using this dataset, we compare the performance of *phyloConverge* with two competing methods that also use strategies to correct for phylogenetic dependence: RERconverge with permulation, as well as the Forward Genomics branch method used by Roscito et al. in their study. For both *phyloConverge* and RERconverge+permulation, we use 500 permulated traits. While *phyloConverge* is capable of producing scores at any resolution, in this analysis, *phyloConverge* scores are computed by fitting the entire CNE sequence to ensure fair comparison with the other methods.

**Figure 2.**
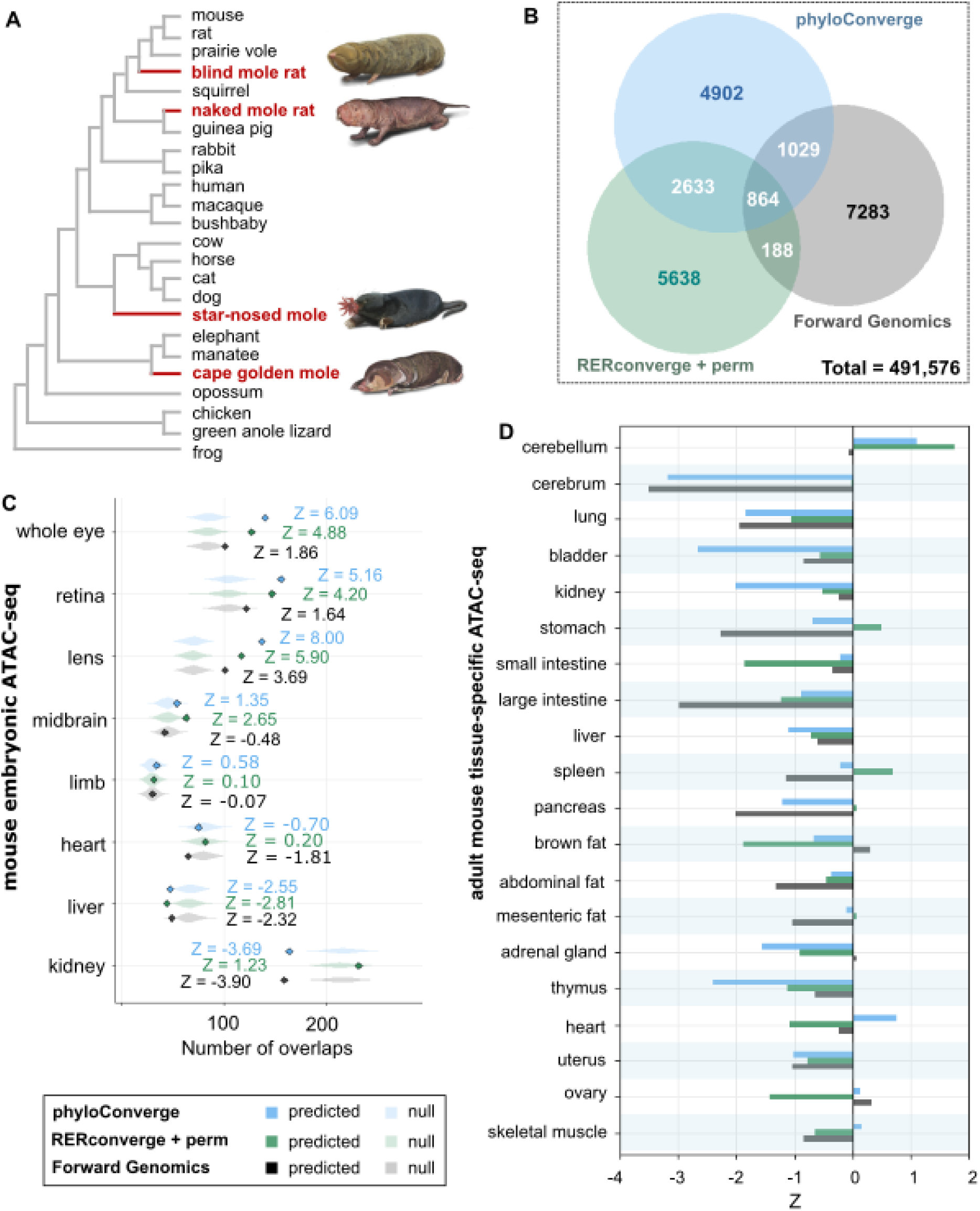
Benchmarking *phyloConverge* with the evolution of conserved non-coding elements (CNEs) in the subterranean mammal phenotype. (**A**) The phylogenetic tree of the benchmarking dataset, containing a multiple genome alignment of 24 species with 4 independent subterranean lineages (red branches) and mouse as the reference genome. (**B**) Venn diagram showing the number of overlaps between top subterranean-accelerated CNEs identified by different comparative methods, *phyloConverge*, Forward Genomics, and RERconverge+permulation. (**C**) Correlation between the top-ranking CNEs identified by the 3 methods with tissue-specific open chromatin regions in mouse embryonic tissues. (**D**) Correlation between the top-ranking CNEs with tissue-specific open chromatin regions in adult mouse tissues (plotted only as Z-scores).

We score the 491,576 CNEs for acceleration using both *phyloConverge* and RERconverge and identify the top-ranking CNEs computed by each method. To evaluate the quality of the rankings produced by different methods fairly, we set permulation p-value thresholds that would approximately match the number of subterranean-accelerated CNEs identified by Roscito et al. with Forward Genomics (9,428 CNEs with p-value ≤ 0.032 for *phyloConverge*, and 9,323 CNEs with p-value ≤ 0.023 for RERconverge). We compare the three size-matched sets of top-ranking CNEs identified by the three methods and find that only 864 CNEs (∼9.2% of each set) are commonly identified by all three methods (Fig. 2B). *phyloConverge* and RERconverge share 3,497 (∼37.3%) commonly identified top-ranking CNEs, which is notably more than those shared between Forward Genomics and *phyloConverge* (1,893 CNEs, ∼20.2%) as well as between Forward Genomics and RERconverge (1,052 CNEs, ∼11.2%). Despite the moderate number of overlaps, we highlight that given the large sample space of 491,576 CNEs, the overlaps between the sets are statistically significant compared to random chance (Z-scores of 256.3 for *phyloConverge* and RERconverge, 132.4 for *phyloConverge* and Forward Genomics, and 67.0 for RERconverge and Forward Genomics). We note that *phyloConverge* and RERconverge are quite different in their model specifications, and that statistical testing procedures with RERconverge and Forward Genomics are more conceptually similar. The main point of similarity between RERconverge and *phyloConverge* is that both rely on maximum likelihood estimates of evolutionary rates and both use the permulation bias correction, suggesting that these features drive the observed overlap.

Next, for each set of top-ranking CNEs, we evaluate the association between the subterranean-accelerated CNEs with tissue-specific open chromatin regions (OCRs), hereby termed “marker OCRs”, across several mouse embryonic tissues (see list of identifiers in Supplementary Table S1). Among the common traits shared by subterranean mammals is that reduced reliance on vision results in degenerated visual structures. These species have small eyes and are either effectively blind or have very low vision capacity (Sweet 1909; Sanyal et al. 1990; Catania 1999; Hetling et al. 2005). As such, we expect that eye-related regulatory regions would experience relaxed selection and exhibit greater divergence. By computing the number of intersections between subterranean-accelerated CNEs and marker OCRs, we observe that all three methods produced sets of subterranean-accelerated CNEs with strong enrichments for the marker OCRs of the whole eye, retina, and lens, relative to 1,000 randomly selected size-matched null sets of CNEs (Fig. 2C), which agrees with our expectation. Notably, the enrichments for eye-related regions are the strongest for *phyloConverge* compared to competing methods. Meanwhile, no enrichment is observed for the marker OCRs of the control non-ocular tissues, including limb, heart, liver, and kidney, for which an enrichment is not expected.

Interestingly, *phyloConverge* and RERconverge produce correlation with midbrain marker OCRs, which is not detected by Forward Genomics. This signal is consistent with the observation that specific midbrain structures that receive direct optical input (superior colliculus and lateral geniculate nucleus) are highly atrophied in subterranean mammals, although the sizes of most structures in the midbrain are comparable to mice (Cooper et al. 1993; Crish et al. 2006). A similar effect is observed in the cave-dwelling Pachón ecotype of the Mexican tetra fish, which possesses degenerated visual structures (Moran et al. 2015). These observations lend support to the hypothesis that eye degeneration is accompanied by complementary changes in brain structure that is detectable as reduced constraint on some brain-specific regulatory regions.

We perform additional enrichment analysis on an atlas of tissue-specific chromatin accessibility across 20 adult mouse tissues (Liu et al. 2019). We firstly note that the enrichment signals for the embryonic tissues (Fig. 2C) are generally stronger than those for the adult tissues (Fig. 2D), consistent with the findings that many deeply conserved enhancers function as drivers of developmental processes (Plessy et al. 2005; Dickel et al. 2018; Yuan et al. 2018). While most of the control tissues are depleted of top-ranking subterranean-accelerated CNEs, *phyloConverge*- and RERconverge-identified top-ranking regions are enriched for marker OCRs of the adult cerebellum (Fig. 2D). The cerebellum is a major structure in the hindbrain that regulates motor coordination (Reeber et al. 2013), cognitive and emotional processing (Schmahmann 2010), as well as ocular motor control (Kheradmand and Zee 2011). In naked mole rats, the cerebellar region involved in visual signal processing has indeed been reported to be degenerated, coinciding with the expansion of the region for the somatosensory system (Marzban et al. 2011) that facilitates the processing of tactile cues for navigation (Sarko 2013). Overall, these results demonstrate that *phyloConverge* is able to improve the prediction of genomic elements with high specificity of phenotypic associations, compared to competing methods.

### Phylogeny-aware calibration is important for avoiding false positives due to confounders

One of the challenges of using comparative genomics to identify regions associated with a specific phenotype is that the phenotype-unaware conservation signal is already highly associated with functional data. Thus, we evaluate whether statistical calibration with permulation can improve the specificity of prediction. First, we evaluate the correlation between the global scaling factor *ρ_o_* (equivalent to computing the phylogeny-wide conservation score) and the corrected scores (*s_corr_*) computed by *phyloConverge*, versus the uncorrected scores (*s_uncorr_*) computed by *phyloP*. Without bias correction, there is indeed a negative correlation between *ρ_o_* and the magnitude of convergent rate shift (Fig. 3A, top). This means that without statistical calibration, stronger convergent signals are given to elements that are more strongly conserved. *phyloConverge* calibrates for this covariate (Fig. 3A, bottom). Repeating this experiment with CNE length as another covariate, we find a similar trend with longer regions being more likely to produce strong convergent rate shift scores (Fig. 3B). Again, *phyloConverge* corrects for this bias. All in all, these observations suggest that permulation can improve prediction specificity by correcting for potential confounders.

**Figure 3.**
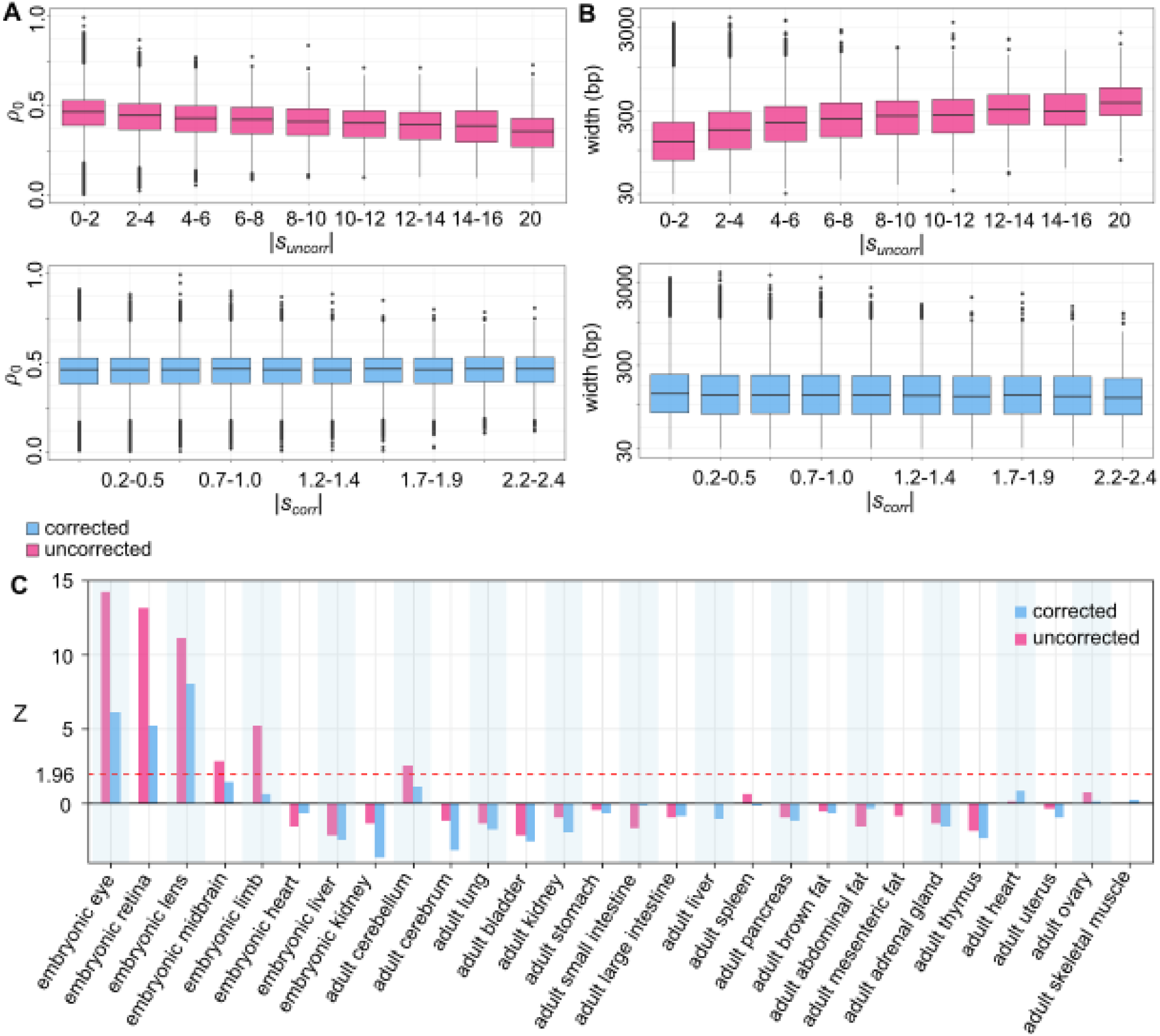
Phylogeny-aware calibration of test statistics improves prediction specificity and bias reduction. (**A**) Without bias correction, the magnitude of the conservation/acceleration score |*s_uncorr_*| is significantly negatively correlated with phylogeny-wide conservation signal *ρ_o_* (top). With bias correction, this correlation is corrected (bottom). (**B**) Without bias correction, the |*s_uncorr_*| has a significant positive correlation with element width, while calibration corrects for this bias. (**C**) Top-accelerated CNEs identified with calibration show improved specificity of enrichment for expected functions.

In order to quantify how much the presence and removal of this statistical bias affects biological inference, we evaluate the functional associations of the top-ranking accelerated elements that are identified with bias correction versus without. We set the threshold for the uncorrected scores *s_uncorr_* at -5.1 to obtain an approximately size-matched set of top-ranking accelerated CNEs (resulting in 9,325 CNEs) and, as before, correlate their coordinates with marker OCRs of mouse tissues (Fig. 3C). The results show that top-ranking accelerated CNEs identified without correction show very strong associations (way above the 95% confidence Z-score of 1.96, red dashed line) with embryonic tissues, including tissues for which strong relaxation of genetic elements is not expected such as limb and midbrain. Strong associations are also observed for the adult cerebellum, which would suggest a pervasive deterioration of the organ. Applying bias correction dramatically improves the tissue specificity of the acceleration signal, highlighting eye and neuronal tissues as expected. Notably, we observe that accelerated regions are enriched for parts of the brain that are relatively constrained in connectivity and function (such as the midbrain and cerebellum), and strongly depleted for cerebrum, which is plastic.

### Subterranean-accelerated elements are enriched for ocular and neuronal functions

To evaluate the functional associations of the top subterranean-accelerated CNEs identified by *phyloConverge* in detail, we use the Genomic Regions Enrichment of Annotations Tool (GREAT) (McLean et al. 2010) to associate the 9,428 CNEs with genes based on distance, and compute CNE enrichments for each gene (see Methods). We find that 76 out of 21,395 genes in the Ensembl database are significantly enriched for the subterranean-accelerated CNEs [hypergeometric test false discovery rate (FDR) ≤ 0.05] (Supplementary Table S2). We then evaluate the enrichment for canonical pathways among these 76 genes. Out of 1,330 canonical pathways, we identify 9 top-ranking pathways (enrichment p-value ≤ 0.05) (Fig. 4A). Notably, these pathways regulate interrelated processes relevant to ocular and/or neuronal functions. For example, the calcium/calmodulin-dependent (Ca-CaM) protein kinase activation pathway plays a role in photoreceptor-regulated light adaptation and maintains the circadian rhythmicity of the mammalian retina (Ko 2020). One of the genes in the Ca-CaM pathway around which significant distribution of subterranean-accelerated CNEs are found, *CAMKIID*, has been found to regulate choroidal and retinal neovascularization in mice (Ashraf et al. 2019). The peroxisome proliferator-activated receptor γ coactivator-1 α (PGC1α) pathway regulates energy metabolism in photoreceptors and similarly manages light susceptibility (Egger et al. 2012), retinal angiogenesis (Saint-Geniez et al. 2013), and circadian clock (Liu et al. 2007). Finally, the paired-like homeodomain transcription factor 2 (Pitx2) pathway is controlled by the Wnt/nuclear β-catenin pathway, and in turn mediates the downstream effects of the Wnt pathway in stabilizing mRNAs (Briata et al. 2003). *PITX2* is critical for eye morphogenesis, and mutations in *PITX2* can cause eye defects and neurodegeneration (Chen and Gage 2016).

**Figure 4.**
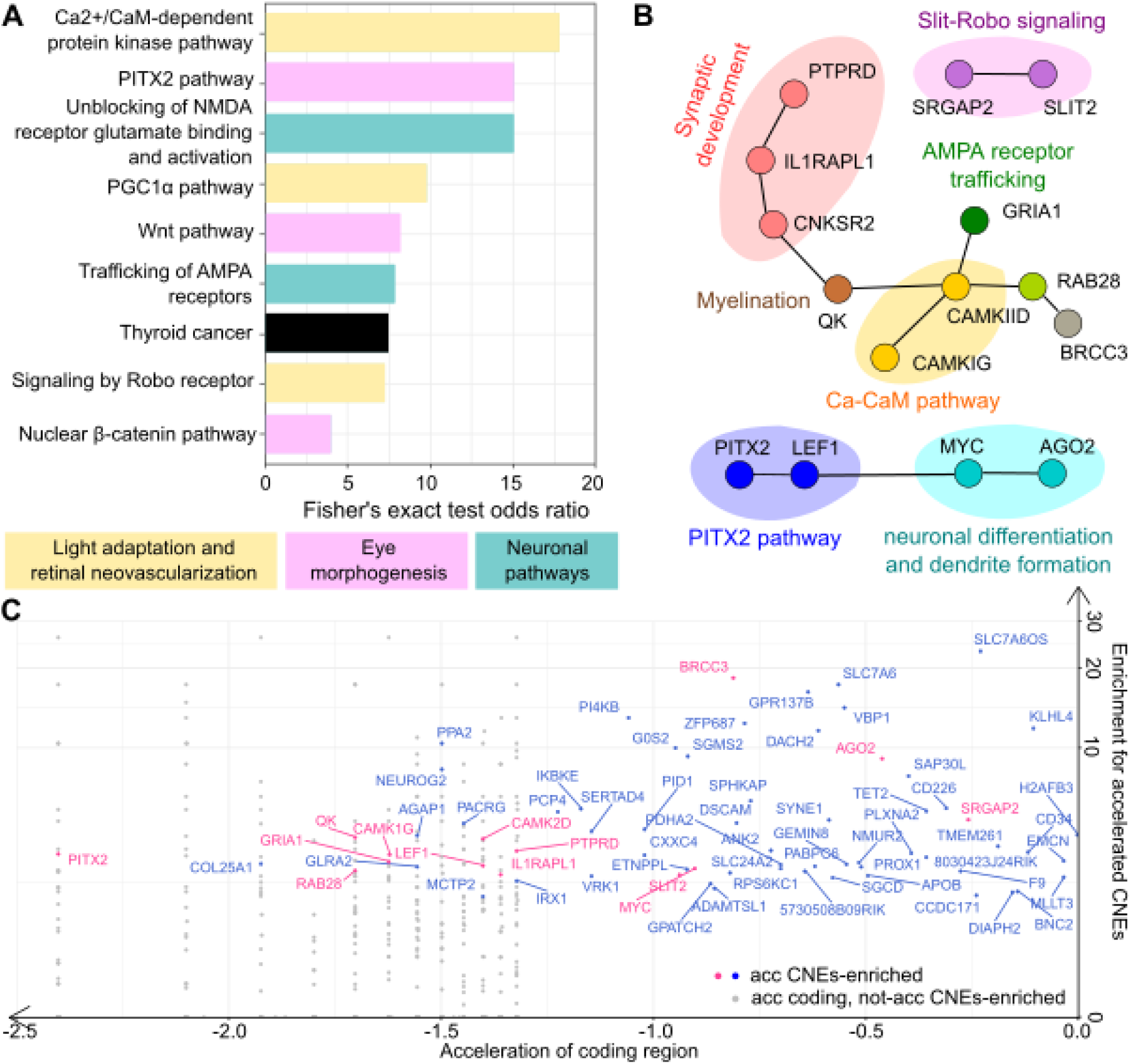
*phyloConverge* identifies subterranean-accelerated CNEs that are distributed around genes associated with ocular and neuronal functions. (**A**) Top-ranking canonical pathways that are enriched for subterranean-accelerated CNEs. (**B**) Protein-protein interactions among some genes that are significantly enriched for subterranean-accelerated CNEs. (**C**) Genes that are enriched with accelerated CNEs (blue and pink dots) can have varying evolutionary rate acceleration in the protein-coding regions, while some genes can have strongly accelerated coding regions without enrichment for accelerated CNEs (grey dots). Note: negative score (*s_corr_*) denotes stronger acceleration.

To further characterize the interactions between the top-enriched genes, we used the Search Tool for the Retrieval of Interacting Genes/Proteins (STRING) (von Mering 2004) to identify significant protein-protein interactions (PPIs) in the set, computing statistical confidence based on available information on gene co-expression, co-occurrence, fusion, proximity, and protein homology. We find that there is an enrichment of PPIs among the top-enriched genes (p-value 0.0179). The identified interactions also reveal additional functional associations (Fig. 4B) that are not captured in pathway enrichment analysis. For example, *CAMKIID* is co-expressed with *GRIA1*, which encodes for glutamate ionotropic receptor AMPA Type subunit 1 (GluA1). GluA1 is a subunit of AMPA receptors, which is involved in facilitating excitatory synaptic transmission and regulates synaptic plasticity in the brain. Indeed, the first cytoplasmic domain of GluA1 contains a residue S567 that serves as a substrate for CAMKII in the regulation of AMPA receptor trafficking (Lu et al. 2010). STRING also identifies interaction between *CAMKIID* and *RAB28*, a RAS-related small GTPase that is highly expressed in photoreceptors and has been associated with cone-rod dystrophies (Iarossi et al. 2020), as well as genes involved in myelination (*QK1* (Wu et al. 2002)) and synaptic development (*CNKSR2* (Zhang et al. 2020), *IL1RAP1* and *PTPRD* (Yamagata et al. 2015)).

Finally, STRING highlights other sets of interacting proteins that further emphasize the tight conjunction between ocular and neuronal functions. These include the co-expression of the Slit-Robo GTPase-activating protein encoded by *SRGAP2* (regulates dendritic spineogenesis and neurite growth (Lucas and Hardin 2017)) with *SLIT2* (required for retinal angiogenesis (Rama et al. 2015)), as well as the interaction between the Pitx2 pathway and proteins that regulate neuronal differentiation and dendrite formation (*MYC* (Zinin et al. 2014), *AGO2* (Rajgor et al. 2018)). All in all, these observations imply that the ocular degeneration resulting from subterranean adaptation may concurrently cause degeneration or remodeling of other nervous system elements.

### Genomic distribution of accelerated CNEs reveals insights into the pleiotropy of regulated genes

To further understand the functional implications of CNE degeneration in subterranean mammals, we use *phyloConverge* to score the acceleration of 19,816 protein-coding regions genome-wide, and evaluate each gene on its coding region acceleration as well as its enrichment for accelerated nearby CNEs. We observe that some genes show strong coding region acceleration but no enrichment for accelerated CNEs, whereas genes that show strong enrichment for accelerated CNEs may have different levels of coding region acceleration (Fig. 4C). The gene that shows the strongest acceleration of coding region and significant enrichment for accelerated CNEs is *PITX2*, followed by *COL25A1* (collagen type XXV alpha 1), whose mutation underlies the aberrant oculomotor neuronal phenotype in congenital cranial dysinnervation disorder (Shinwari et al. 2015). Most of the genes shown in Fig. 4B also exhibits moderate or strong acceleration of coding regions (pink dots).

On the other end of the spectrum, there is a subset of genes that are significantly enriched with accelerated CNEs but are not strongly accelerated in the coding regions. The lack of concordance between protein-coding acceleration and enrichment for accelerated CNEs for this set of genes may reflect the role of CNEs in regulating the expression of pleiotropic genes that experience relaxed selection on certain functions but are otherwise still critical for survival. Thus, their protein-coding portions did not accelerate and remain under constraint, but their regulatory elements specific to the vision functions could be under relaxed constraint (acceleration). The top-enriched genes in this set include genes that encode for amino acid transporters (*SLC7A6*), probable nuclear localization of RNA polymerase II (*SLC7A6OS*), regulators of DNA damage response (*BRCC3* and *G0S2*), and a chaperone protein for the Von Hippel-Lindau tumor suppressor gene product (*VBP1*). These functions are general cellular processes that are involved across many tissues, and thus strong conservation of their protein sequences (but not necessarily regulatory elements) would be expected. Examples of eye-related pleiotropy in the set include *DIAPH2*, which has been associated with both age-related macular degeneration and ovarian development (Vladan et al. 2013), and *PROX1*, which is involved in the development of not only the lens and the central nervous system, but also the liver, pancreas, and heart (Oliver et al. 1993; Wigle et al. 1999; Sosa-Pineda et al. 2000; Burke and Oliver 2002; Lavado and Oliver 2007). For these pleiotropic genes, changes that drive the convergent phenotype may have occurred in the regulatory elements that control their expression.

To summarize these trends at the pathway-level, we examine the contrast between pathway enrichment in genes accelerated in coding regions and genes accelerated in CNEs (Fig. 5A). We find that the top-ranking pathways with the strongest enrichment for accelerated coding regions (denoted in blue) are specific to phototransduction, while the pathways enriched for genes with non-coding acceleration point to diverse developmental and neuronal processes and importantly are not enriched for coding accelerations. We provide detailed network views of the top pathways enriched for both acceleration types in Supplementary Figs. S1 and S2.

**Figure 5.**
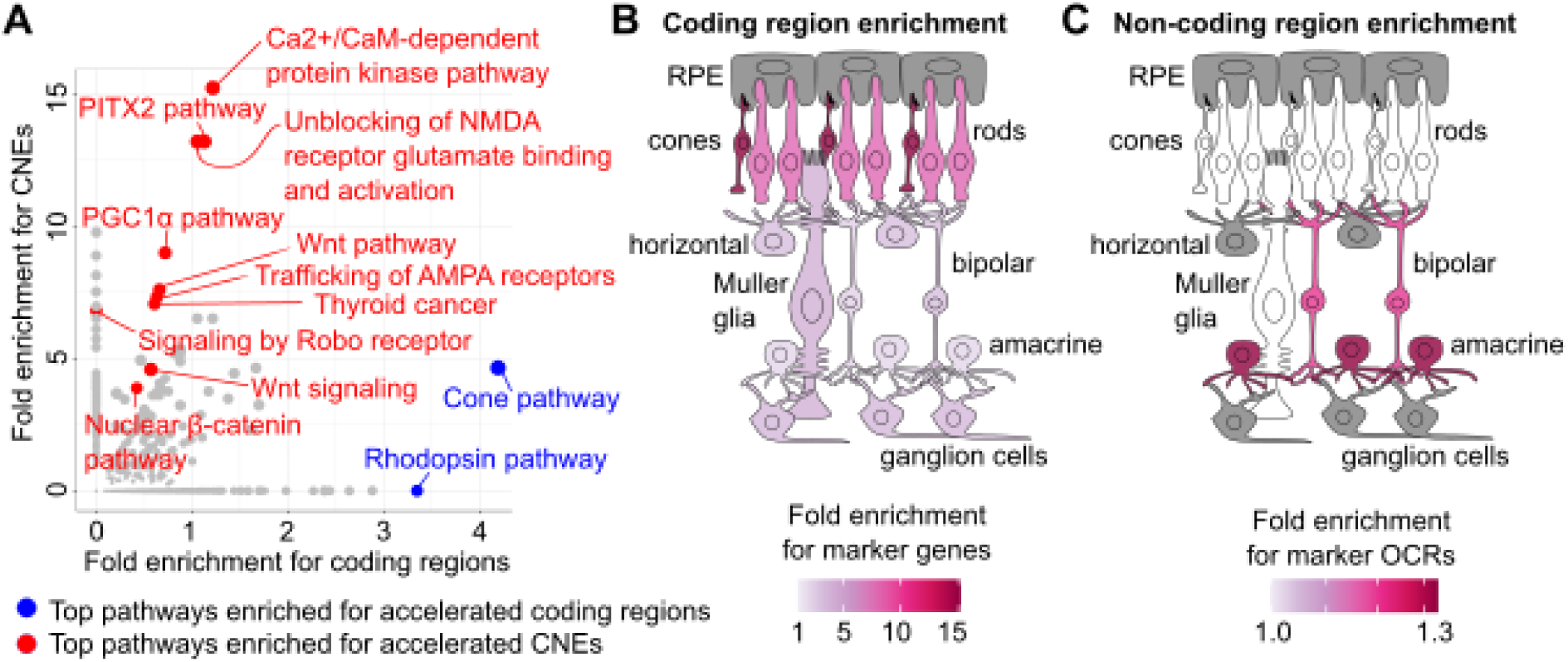
Protein-coding region versus CNE acceleration occurs across distinct biological functions. (**A**) The top-ranking canonical pathways enriched for subterranean-accelerated protein-coding regions (blue) or CNEs (red). (**B**) Genes that are strongly accelerated in the coding regions show the strongest enrichment for cone and rod photoreceptor markers genes. (**C**) Strong enrichments for subterranean-accelerated CNEs are found in retinal cells in the inner nuclear layer, namely the amacrine and bipolar cells. Tissues in (**B**) and (**C**) for which the Benjamini Hochberg adjusted enrichment p-value > 0.05 are colored in white, while tissues for which genetic or genomic annotations are not available are colored in grey.

A similar trend emerges when we examine enrichment for retinal cell-type-specific marker genes and marker OCRs from single cell sequencing experiments (Fig. 5B). Using a curated dataset of tissue-specific marker genes across different retinal cell types (Macosko et al. 2015), we observe that genes with accelerated coding regions are significantly enriched for all retinal cell types, but the cone and rod photoreceptors show drastically stronger fold-enrichment than other tissues (Fig. 5B). In contrast, using genomic annotations of marker OCRs in five retinal cell types (Norrie et al. 2019), we find that the photoreceptor layer and the Müller glia exhibit no enrichment, while cells in the inner nuclear layer of the retina (the amacrine and bipolar cells) show strong, statistically significant enrichment (Fig. 5C). While photoreceptors, amacrine, and bipolar cells are all neuronal cell-types, the photoreceptors are highly specialized for phototransduction, while amacrine and bipolar cells are specialized inter-neurons that perform signal transduction and signal processing function. Bipolar cells are solely responsible for relaying information from the photoreceptors to the inner layers of the retina and performing specific transformation of neuronal signals (Euler et al. 2014), while amacrine cells relay signals from the bipolar cells to the ganglion cells and control the temporal regulation of visual signals (Taylor and Smith 2012). The observation of weak enrichment for accelerated coding regions and strong enrichment for accelerated CNEs in the amacrine and bipolar cells lends further evidence to the argument that subterranean adaptation is accompanied by transformation of neuronal functions, which are driven by changes in transcriptional control rather than the genes themselves.

In summary, using pathway and marker enrichment analyses, we find that coding region and CNE acceleration is concentrated in distinct biological functions. While coding region acceleration is observed in genes whose functions are specific to visual signal transduction, non-coding acceleration is enriched for a broader set of developmental and neuronal processes. These observations support the hypothesis that relaxation of selection in the coding regions is concentrated in highly specialized genes, while pleiotropic genes that contribute to the development and function of the visual system but have additional non-vision related roles experience mostly non-coding relaxation.

### High-resolution scoring reveals acceleration of transcription factor binding motifs relevant to ocular and neuronal development

The general framework of *phyloConverge* has the capacity to fit the convergent rate shifts model at arbitrary small, even base-pair, resolution, which allows for a deeper inquiry into the information content of different parts of a given CNE. To understand how CNE level acceleration is reflected in the TFBS profiles, we identify HOCOMOCO TFBS matches (see Methods) within the subset of CNEs that show large-scale subterranean acceleration. Specifically, we compute the *phyloConverge* acceleration scores specifically on the TFBS matches, generating a site-specific *s_corr_* for each TFBS match. We then calculate the mean *s_corr_* for the given TF motif by averaging the computed site-specific *s_corr_* values, thus generating a global subterranean *TFBS acceleration*. An alternative approach for computing the TF specificity of the acceleration signal would be to simply compute the enrichment of TFBS occurrence in our subterranean-accelerated CNEs, which does not require any local scoring. This is indeed the common method for associating TFs with sets of genomic regions and is widely used in analyzing genomic regions derived from functional data (Ronzio et al. 2020; Yan et al. 2020). We refer to it as *TFBS enrichment* (Fig. 6A).

**Figure 6.**
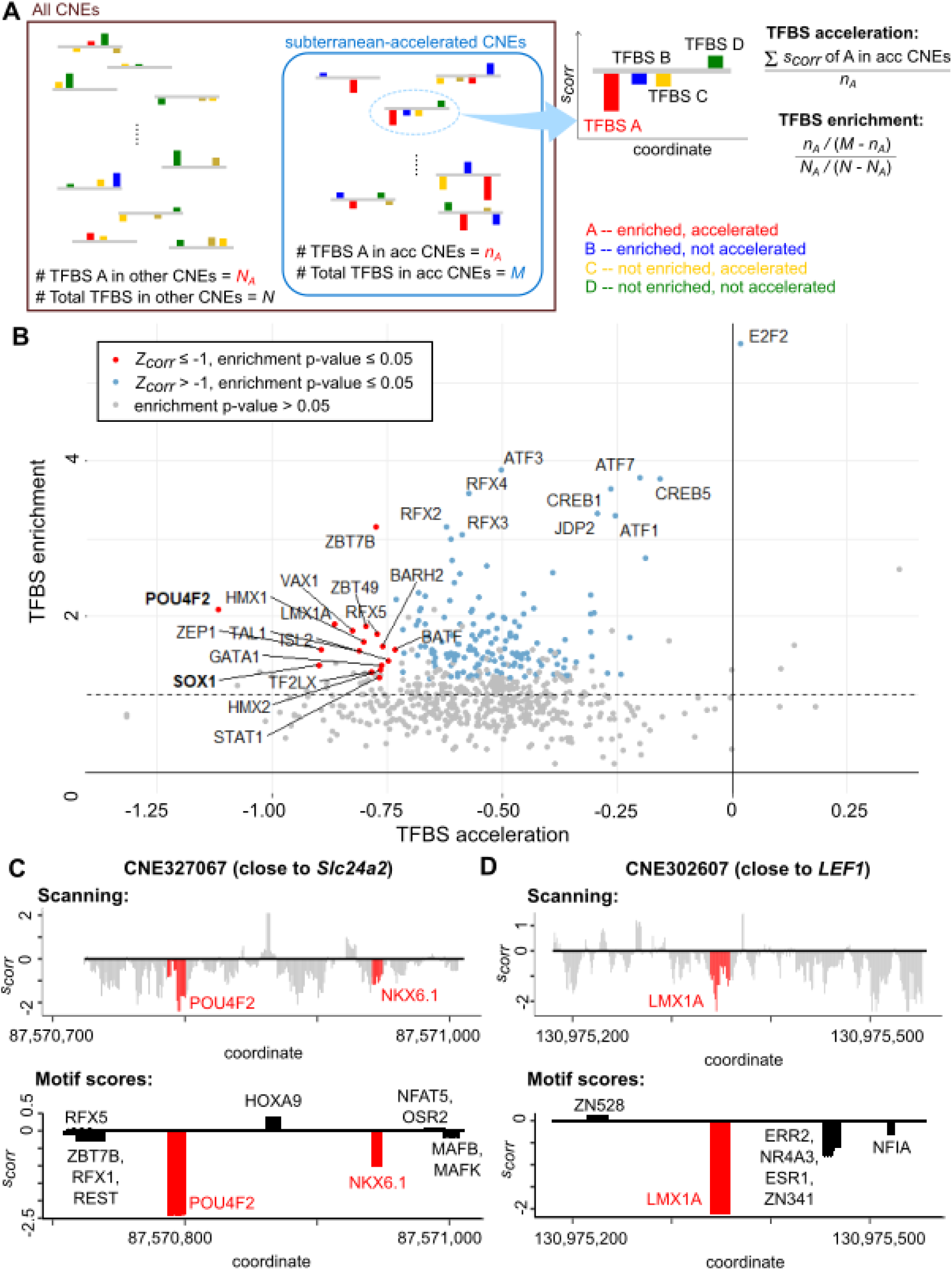
*phyloConverge* can highlight segments of a conserved non-coding element (CNE) that experience strong selection. (**A**) Each transcription factor binding site (TFBS) motif is scored for its average acceleration/conservation score (*s_corr_*) and enrichment for subterranean-accelerated CNEs. For each TFBS motif, genome-wide calls are made and the coordinates that overlap the subterranean-accelerated CNEs are identified. Average *s_corr_* is computed from the coordinates that overlap the subterranean-accelerated CNEs. Enrichment is computed using a hypergeometric distribution. (**B**) While most TFBS motifs are not significantly enriched for subterranean-accelerated CNEs (grey dots), a handful of motifs are (red and blue dots). Some of the enriched motifs also show a stronger acceleration on aggregate (red dots), which include motifs that have been strongly implicated in ocular and neuronal functions. The red dots are identified by converting the mean *s_corr_* into Z-scores (*Z_corr_*), and then setting a threshold of -1. (**C**) High-resolution scanning of CNE327067. In the top panel, each nucleotide is scored by using a window of ±5bp around the nucleotide. The bottom panel shows the TFBS-scale scores of the TFBS motifs in the CNE. (**D**) High-resolution scanning of CNE302607.

Plotting TFBS acceleration against enrichment (Fig. 6B), we find that the two scores are correlated, but not equivalent. The strongest enrichments are observed for the ATF/CREB and the RFX transcription factor families. Members of these families are ubiquitously expressed and have diverse functions in regulating cell growth, inflammation, and apoptosis (Persengiev 2003; Wen et al. 2010; Cui et al. 2021). We also identify a subset of TFBS motifs that exhibit both strong enrichments in subterranean-accelerated CNEs and strong acceleration scores on aggregate (red dots in Fig. 5B) (threshold for strong acceleration is set by converting the mean *s_corr_* values into Z-scores and selecting the motifs with Z-scores ≤ -1). Notably, the top-ranking motifs in this set have been associated with neuronal and/or eye-related functions and developmental processes (see Supplementary Table S3 for the specific roles of each motif). The most strongly accelerated motif that also has strong enrichment is a canonical retinal marker *POU4F2* (POU domain class 4, transcription factor 2), whose expression together with *ISL1* has been found to be sufficient for specifying the retinal ganglion cell fate (Wu et al. 2015). *SOX1* is also a known regulator of neural cell fate determination (Kan et al. 2004) that is critical for lens development (Nishiguchi et al. 1998). This suggests that the relaxed selection for eye and neuronal functions in the subterranean environment causes a widespread divergence of TFBS motifs relevant to those functions.

High-resolution scoring can also segment a given subterranean-accelerated CNE into regions with differing relevance to the convergent phenotype of interest. We illustrate this using two example subterranean-accelerated CNEs shown in Figs. 6C and 6D. CNE327067 (Fig. 6C) is intergenically located in between *Slc24a2* [a cation/calcium ion exchanger that maintains the homeostasis of sodium, potassium, and calcium ion levels in the brain, retinal ganglion cells, and the retinal cone photoreceptors (Sharon et al. 2002)] and *MLLT3* [a regulator of the self-renewal of haematopoietic stem cells (HSCs) (Calvanese et al. 2019)]. We use *phyloConverge* to scan each nucleotide in the CNE and compute the score from a window of ±5bp around the nucleotide (top of Fig. 6C), considering that ∼10bp is the approximate scale of TFBS motifs. We find that the CNE contains strongly accelerated and decelerated segments that correspond to known motifs for transcription factor binding. Strong deceleration is observed for a segment that corresponds to the binding site for *HOXA9* (Homeobox protein Hox-A9), which like *MLLT3* is also a regulator of HSC self-renewal (Ferrell et al. 2005). The coding region of *MLLT3* is in fact also not accelerated (Fig. 4C), overall suggesting a possible functional association between the conserved *HOXA9* motif and *MLLT3*. Meanwhile, the most strongly accelerated segment of the CNE corresponds to the binding site for *POU4F2*. The co-expression of *POU4F2* and *Slc24a2* in retina (Brooks et al. 2019), and the presence of strongly accelerated binding motif for *POU4F2* in this CNE, suggest the possible involvement of *POU4F2* in regulating the transcription of *Slc24a2* specifically during eye development. The CNE also contains a moderately accelerated segment that corresponds to the binding site for *Nkx6.1*, a homeodomain protein that is responsible for determining neural fates and regional patterning (Sander et al. 2000). Together, all these observations suggest that CNE327067 may have a pleiotropic function in regulating the expression of its two closest genes, which underlie different phenotypes that are differentially selected by the subterranean environment.

A second example is illustrated for CNE302607 (Fig. 6D), which is located upstream of *LEF1*, an effector of Wnt signaling that mediates the regulation of *PITX2* (Abu-Elmagd et al. 2010). We similarly scan each nucleotide in the CNE using a window size of ∼10bp and find strong acceleration for a segment that corresponds to the binding site for *LMX1A* (LIM homeobox transcription factor 1 alpha), a midbrain-specific transcription factor (Uhde et al. 2010). *WNT1* and *LMX1A* regulate the midbrain dopaminergic neuronal differentiation via an autoregulatory loop (Chung et al. 2009), which is also mediated by *LEF1* (Nouri et al. 2020). The promoter of *LMX1A* contains a binding site for *LEF1*/T-cell factor (Nouri et al. 2020), but the presence of the strongly accelerated binding site for *LMX1A* in this CNE suggests that a possible feedforward mechanism may be involved in maintaining the levels of *LMX1A* and *LEF1* during development. All in all, these examples demonstrate that *phyloConverge* is able to highlight portions of an element that are potentially important for the convergent phenotype of interest and that then lead to hypotheses that are amenable to experimental interrogation.

## Discussion

We introduce *phyloConverge*, a new comparative genomics method that combines explicit estimation of nucleotide substitution rates and adaptive calibration of test statistics to identify the phenotypic associations of genetic elements. For a phenotype of interest, *phyloConverge* quantifies the amount of local rate convergence signal via a maximum likelihood estimation of a two-parameter neutral tree scaling model. The MLE statistics are calibrated with an empirical p-value, which dramatically reduces multiple sources of bias.

Benchmarking our method using an empirical dataset that was previously analyzed for rate convergence in subterranean mammals (Roscito et al. 2018), we find that *phyloConverge* identifies CNEs that exhibit strong associations with ocular functions–which satisfies our expectations for the phenotype– and discover that the regression of ocular functions may be accompanied by changes in neuronal functions and development. We also demonstrate that *phyloConverge* can analyze a given CNE in segments and provide insights about its pleiotropic activity in the specific phenotypic context. Importantly, *phyloConverge* produces unbiased signals because it corrects for biological and technical confounders.

*phyloConverge* offers a scalability to perform rapid, calibrated scoring at flexible resolution. We have demonstrated this flexibility by applying *phyloConverge* in three complementary ways: scoring entire CNEs for aggregate CNE-level acceleration, scoring TFBS for TFBS-level acceleration, and dissecting aggregate acceleration signals with high resolution scoring. This highly flexible framework allows for rapid convergent acceleration scanning with less computational overhead than competing methods. For example, to score 1 million elements, RERconverge would require the pre-estimation of 1 million element-specific trees, and Forward Genomics would require computations of the neutral tree model as well as local (branch-specific) or global (relative to the root) percent identity values per branch per element. For the same analysis, *phyloConverge* would only require the estimation of one neutral tree model, while the scoring of the 1 million elements would be performed through a small number of parameter estimations and hypothesis testing. While adding the permulation step incurs additional computational cost, we have demonstrated previously the advantage of calibrating the resulting statistics via permutations is not method-dependent.

While in this analysis we have focused on pre-specified CNEs, *phyloConverge* can tractably be extended to perform convergence scans genome-wide, generating convergent rate tracks similar to the *phyloP* conservation score. This provides the option of scanning entire genome alignments to detect coordinates with significant convergent shifts in evolutionary rates without needing prior knowledge about the coordinates and definitions of the functional regulatory units of the elements. However, the optimal pipeline for such unsupervised genome-wide scanning remains to be determined, because we still lack a thorough understanding of non-coding regions to inform our interpretation of significant rate shifts in non-coding elements. For example, in Figs. 6C and D, we scan each nucleotide position in 2 CNEs to find the relative positions of TFBS motif-scale segments that show convergent rate shifts in either direction. While some strongly accelerated segments actually correspond to known TFBS motifs, we also observe strong rate shifts at coordinates where no known binding motifs are found, and their significance is unclear. Furthermore, there are also other types of non-coding elements whose conservation patterns are less well-understood, including long non-coding RNAs, which tend to have a highly conserved promoter region but a less conserved transcribed region (Johnsson et al. 2014), and microRNAs, which can also have varying conservation patterns (Praher et al. 2021). Further investigation into understanding conservation in non-coding elements and how it can inform the design of unsupervised genome-wide scanning for convergent rate shifts can be pursued in future work.

Finally, it is important to note that predictions generated by sequence alignment-based methods such as *phyloConverge* should be interpreted with some caveats. It is increasingly understood that some enhancers can have consistent functional activity across distantly related species despite lacking enhancer-wide sequence conservation. For example, characterization of the putative *Islet-Spacer* enhancers in sponge, fish, mouse, and human revealed that functionally homologous enhancers can have high variability in compositions, orientations, numbers, and alignments of a common set of TFBS (Wong et al. 2020). The quality of predictions also hinges upon the global alignability of sequences, which can deteriorate with increasing evolutionary distance. For such enhancers, sequence alignment-based methods would likely fail. In this instance, “alignment-free” methods that compare sequences in some functional readout space may be appropriate. Nonetheless, sequence alignment-based methods would be sufficiently powerful to analyze both promoter regions and strongly conserved enhancers that are often critical for developmental processes.

## Methods

### phyloConverge

*phyloConverge* computes the association between convergent genetic changes and convergent phenotypic adaptation using a combination of maximum likelihood-based estimation of evolutionary rate shifts and phylogenetic statistical calibration of test statistics. The input of *phyloConverge* includes a multiple sequence alignment (MSA), a phylogenetic model of neutral nucleotide substitution (which can be estimated from sites that are expected to undergo neutral evolution, e.g., fourfold-degenerate sites), and the list of species with the convergent phenotype. To quantify the uncorrected association score between the evolutionary rate of a genetic element and the convergent phenotype, *phyloConverge* uses the *phyloP* function in the *RPHAST* package (Pollard et al. 2010; Hubisz et al. 2011). Given the neutral nucleotide substitution rates and branch lengths in the phylogenetic model computed from neutral sites, *phyloP* performs maximum likelihood estimation to scale the branch lengths in the model to optimize the probability of observing the given MSA. The null hypothesis–that the foreground (convergent) branches are not differentially scaled from the rest of the branches–requires the estimation of one phylogeny-wide branch scaling factor *ρ_o_*. The alternative hypothesis–that the foreground branches are differentially scaled from the rest of the branches–requires the estimation of two scaling factors: a phylogeny-wide scaling factor *ρ_1_* and a foreground-specific scaling factor *λ*. *phyloP* then performs a hypothesis test to determine whether the null model or the alternative model better fits the observed MSA. *phyloConverge* uses the likelihood ratio test (LRT) for the hypothesis testing and adopts the ‘CONACC’ scoring method defined by *phyloP* (positive scores denote conservation, negative scores denote acceleration).

To empirically calibrate for statistical biases, *phyloConverge* performs permutation tests by producing numerous null or ‘fake’ phenotypes and using *phyloP* to compute null scores from them. To generate the null phenotypes, we previously developed a phylogeny-aware strategy for permuting phenotype trees that is grounded on phylogenetic simulation, called permulation (Saputra et al. 2021). The objective of the permulation strategy is to generate permutations that capture the existing structure in the data. Permulation achieves this by preserving the phylogenetic features in the observed phenotype, which in binary traits are captured by the number of foreground species and the phylogenetic dependence between the foreground species. First, the method performs a Brownian motion phylogenetic simulation based on the given trait tree and neutral tree. Next, the simulated branch length values are used to inform the selection of a new set of branches that will be proposed as a candidate null phenotype, specifically by choosing the branches with top-ranking branch length values. A rejection sampling process is then employed to determine if the candidate null phenotype would be accepted as valid, in which two conditions have to be met: that the null phenotype has the same number of foreground branches, and that the foreground branches have matching phylogenetic relationship as observed in the true phenotype (Fig. 1B). After numerous valid null phenotypes are obtained, *phyloP* is used to compute scores for each of the null phenotypes, such that a null distribution of *phyloP* scores for the given MSA is obtained. We previously developed two permulation strategies for binary phenotypes: the ‘complete case’ (CC) method, which produces null phenotype trees from the complete topology, and the ‘species subset match’ (SSM) method, which accounts for missing sequences in a particular MSA. While the SSM method is more stable and accurate, the CC method is significantly faster, with comparable if slightly less accuracy. For tractability, *phyloConverge* currently makes use of the CC method.

In such a permutation test, the significance of deviations from the expected value is typically measured by computing empirical p-values that are defined as the proportion of the null statistics that are as extreme or more extreme than the observed test statistic. In a two-tailed test, these extreme values make up the area under the curve beyond the observed statistic and the negative of the observed statistic. As *phyloP* defines acceleration as a negative score and conservation as a positive score following the ‘CONACC’ scoring mode, the two tails of the null score distribution signify opposing directions of rate shift, where the lower tail denotes acceleration and the upper tail denotes deceleration. Because of this directionality and because the null distribution is not necessarily trivial or symmetric (e.g., histogram in Fig. 1A), we calculated the two-tailed empirical p-value *p_corr_* using the two-sided conditional p-value approach described by Kulinskaya (Kulinskaya 2008), which transforms one-sided p-values into equivalent, weighted two-sided p-values for symmetric or asymmetric distributions. Suppose the distribution of null *phyloP* scores follows a strictly increasing continuous cumulative distribution function *F*. Then, *p_corr_* is computed as follows:

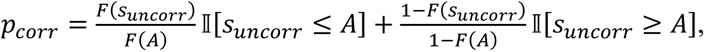

where *s_uncorr_* is the uncorrected score (computed by *phyloP*) for the observed phenotype and *A* is the value that the null distribution is centered on. For our purposes, *A* was chosen as the median of the null scores, such that the weights at both the left and right sides of *A* were equal. Subsequently, the bias-corrected conservation/acceleration score *s_corr_* is computed as the negative logarithm of *p_corr_*, signed by the relative position of *s_uncorr_* with respect to *A*, as follows:

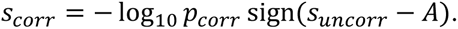

Finally, to improve the computational tractability of permulations, we incorporated a simple strategy to adaptively terminate permulations when a target significance threshold had been reached. For example, suppose we would like to control for significance threshold *α* = 0.05 with a maximum of 1000 permutations. For a genetic element to be significantly associated with the convergent trait, there can only be a maximum of 50 null scores that are as extreme or more extreme than the observed uncorrected score. Formally, suppose we want to control the test for a significance level of *α*, and we set a maximum of *N* permulations. Denoting *S’* as the set of computed null statistics, for a hypothesis to be statistically significant at *α* significance level, the maximum number of null statistics that are as extreme or more extreme than the true statistic, *s_uncorr_*, is therefore *αN*, defined as the “pruning” threshold. At every permulation iteration *i*, we track whether or not the pruning threshold has been reached, given the value of the median of the null distribution at iteration *i*, *A_i_*. If the threshold has been reached, the adaptive *p_corr_* is computed as follows:

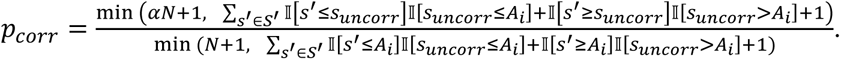

The addition of ‘+1’ to each term is done to correct the tail ends of the distribution. The approach indeed offers remarkable improvements in speed -- parallelizing over 60 cores on one compute node with 95GB memory, the scoring of ∼36,000 CNEs with 500 permulations can be completed in ∼1.5 hours. Using ∼5,000 randomly selected subset of the CNEs dataset and the subterranean foregrounds, the empirical p-values calculated from all 500 permulations (*p_total_*) and the adaptive empirical p-values (*p_corr_*) computed to control significance levels *α* of 0.05 correlate very well (Pearson’s R = 0.978, p-value < 2.22 x 10^-16^), with negligible loss in resolution within the significance level that is controlled (Supplementary Fig. S3). However, we note the necessity for weighing the trade-off between the maximum number of permulations and the *α* level to control for. For example, with a maximum of 500 permulations, setting *α* = 0.01 means that only a maximum of 5 extreme null scores are allowed such that the computations may be prematurely terminated. In such cases, the performance of adaptive permulation may suffer, because stochasticity can cause premature termination of the permulations. The source code for *phyloConverge* is available at GitHub (https://github.com/ECSaputra/phyloConverge).

### Identification of tissue-specific “marker” open chromatin regions (OCRs)

To evaluate the functional enrichments of top-ranking subterranean-accelerated CNEs, we computed the correlations between the CNEs with tissue-specific, “marker” open chromatin regions (OCRs) in mouse tissues. For mouse embryonic tissues, we compiled publicly available ATAC-seq datasets (see Supplementary Table S1 for identifiers). The marker OCRs for the whole eye, retina, and lens were taken directly from Supplementary Data 16 of Roscito et al (Roscito et al. 2018). For the remaining tissues, the datasets with multiple replicates were first pre-processed by identifying consensus regions (regions that were present across at least 2 replicates) using the *GenomicRanges* package in R (Lawrence et al. 2013). Subsequently, the marker OCRs of each given tissue were obtained by subtracting regions that were open in any other tissue from the tissue of interest using BEDTools (Quinlan and Hall 2010).

We also used the chromatin accessibility atlas across adult mouse tissues (Liu et al. 2019). Given that the dataset was presented in the format of consensus peaks, we first identified the OCRs in each tissue by setting the 80^th^ percentile of the read count distribution as a threshold. Then, we identified the marker OCRs of each given tissue by subtracting regions that were open in at least 80% of the other tissues. Finally, the regions were lifted over from the *mm9* to the *mm10* coordinates.

### Benchmarking *phyloConverge* against existing methods

We applied the random subsampling strategy previously used by Roscito et al. (Roscito et al. 2018) to compute correlations between subterranean-accelerated CNEs with marker OCRs. Before computing correlations, we merged nearby subterranean-accelerated CNEs that were within 50bp apart to correct for inflation of significance resulting from multiple CNEs that were very close together. Afterwards, for each tissue, we used BEDTools to find the number of intersections between the marker OCRs of the tissue and the subterranean-accelerated CNEs. We then subsampled 1,000 matched-sized sets of randomly selected CNEs from the total set of CNEs and similarly found the number of intersections with the marker OCRs in order to obtain the null distribution. The strength of correlation between the subterranean-accelerated CNEs and the marker OCRs were quantified as the Z-score computed with respect to the null distribution. This analysis was performed for the top-ranking subterranean-accelerated CNEs from the three methods tested, namely Forward Genomics, *phyloConverge*, and RERconverge with permulations.

To quantify the agreement between two sets of top-ranking subterranean-accelerated CNEs identified by two different comparative methods, we first noted the number of overlapping CNEs between the two sets using BEDtools. Then, we subsampled two sets of randomly selected CNEs from the total set, containing matching numbers of CNEs as the two actual sets, and similarly noted the number of overlapping CNEs. We performed the random subsampling 1,000 times to obtain a null distribution of the number of overlapping CNEs between two randomly selected sets of CNEs with the given sizes. The actual number of overlaps was then converted to a Z-score with respect to the null distribution.

### Pathway enrichment analysis

To associate subterranean-accelerated CNEs with genes, we used the Genomic Regions Enrichment of Annotations Tool (GREAT) (McLean et al. 2010). GREAT associates genes with proximal or distal CNEs using a default association rule called ‘basal-with-extension’. GREAT first determines a ‘basal regulatory region’ around each gene, defined as the window within 1kb downstream and 5kb upstream of the transcription start site (TSS) regardless of overlaps with neighboring genes. Then, the regulatory domain of the gene is extended until it overlaps the basal regulatory region of neighboring genes, up to 1Mb both upstream and downstream. Afterwards, using the set of subterranean-accelerated CNEs as the “foreground regions” and the total CNEs as the “background regions”, GREAT performs hypergeometric tests to compute the enrichments for foreground regions in each gene’s regulatory domain, relative to the superset of background regions. We used GREAT to evaluate the enrichments for 21,395 Ensembl genes, and determined the significance cutoff as Benjamini-Hochberg adjusted p-value ≤ 0.05.

After identifying genes that were significantly enriched for subterranean-accelerated CNEs, we performed enrichment analysis on 1,330 genesets in the canonical pathways, using Fisher’s exact test with the CNE-enriched genes as foreground regions and all the genes across the genesets as backgrounds. The cutoff for top-ranking enriched pathways was set as Fisher’s exact test p-value ≤ 0.05. We also evaluated protein-protein interactions (PPI) between enriched genes using the Search Tool for the Retrieval of Interacting Genes/Proteins (STRING) (von Mering 2004). STRING uses a pooled database combining imported information from existing PPI databases and *de novo*-identified gene co-occurrences, fusions, and neighborhoods as a basis for computing confidence scores of associations for all pairs of proteins. We excluded associations that were determined from text mining, as the associations identified from this approach might be imprecise.

### Enrichment analysis on retinal cell-type-specific marker genes and marker OCRs

We performed enrichment analysis on the top-ranking genes that were subterranean-accelerated in the coding regions using the retinal tissue-specific marker genes produced in Macosko et al. (Macosko et al. 2015) as validation datasets. The same permulation p-value threshold that was used to define the top-ranking subterranean-accelerated CNEs (p-value ≤ 0.032) was also used to define the top-ranking subterranean-accelerated coding regions. To perform enrichment analysis on the top-ranking subterranean-accelerated CNEs, we used a dataset of single cell ATAC-seq regions across different retinal tissues (Norrie et al. 2019). Clustering of single cells was performed using Seurat (Hao et al. 2021) and Signac (Stuart et al. 2021) for the single cell RNA-seq and single cell ATAC-seq data, respectively, and tissue type assignments were made by integrating the multimodal datasets and transferring the single cell RNA-seq cluster labels to the corresponding single cell ATAC-seq clusters (Stuart et al. 2019). Cell-type-specific marker OCRs were finally defined by finding the differentially accessible ATAC-seq peaks for the five resulting clusters (rods, cones, bipolar cells, amacrine cells, Müller glia). For both the coding and non-coding analysis, enrichment analysis was performed using the hypergeometric test. The cutoff for significant enrichment was set as Benjamini-Hochberg adjusted hypergeometric p-value ≤ 0.05.

### Analysis of transcription factor binding site (TFBS) motifs

Genome-wide scanning for possible TFBS motifs was performed using PWMScan (Ambrosini et al. 2018). The parameters for PWMScan include the genome assembly of interest, the position weight matrix (PWM) of the motif of interest, and a threshold cutoff for calling the TFBS motif. We obtained the PWMs of 759 TFBS motifs from the HOCOMOCO database (version 11) (Kulakovskiy et al. 2018). We then used *motifDiverge* (Kostka et al. 2015) to compute the background frequency of each nucleotide based on the probability matrix of a given TFBS, and infer the PWM matrix of each TFBS and the TFBS calling cutoff to use (set to control Type I error rate below 10^-5^). Using these input parameters, we previously made the genome-wide TFBS calls for other uses with the human *hg19* coordinate from the UCSC Genome Browser (Kent et al. 2002). These genome-wide calls were then lifted over to the mouse *mm10* coordinate using the liftOver tool from UCSC (Kuhn et al. 2013).

Given the TFBS calls, we computed two metrics for each TFBS motif: TFBS acceleration and TFBS enrichment (Fig. 6A). We first identified the overlapping coordinates of a given motif among subterranean-accelerated CNEs using the *intersect* function of BEDtools. Each of these overlapping coordinates were then scored with *phyloConverge* using 500 maximum permulations, and the average *s_corr_* from these scoring were computed. With reference to Fig. 6A, to calculate motif enrichment, we used a hypergeometric test to compute the probability and fold-enrichment of finding *n_A_* overlapping coordinates of motif *A* out of *M* total overlapping coordinates of all motifs, given that there were *N_A_* + *n_A_* total coordinates of motif *A* genome-wide.

## Data Access

The multiple genome alignment dataset and CNEs used in this article has been previously published in Roscito et al. (Roscito et al. 2018) and are available at https://bds.mpi-cbg.de/hillerlab/CNEDivergence/.

All publicly available datasets used as validation data are listed in Supplementary Table S1.

## Acknowledgments

This work was supported by the National Institutes of Health (R01 HG009299 to N.C. and M.C.).

